# Overexpression of *bla*_OXA-58_ Gene Driven by IS*Aba3* is Associated with Imipenem Resistance in a Clinical *Acinetobacter baumannii* Isolate from Vietnam

**DOI:** 10.1101/2020.06.29.178632

**Authors:** Anh T. Nguyen, Son C. Pham, Anh K. Ly, Chau V.V. Nguyen, Thanh T. Vu, Tuan M. Ha

## Abstract

The aim of this study was to investigate genetic structures and expression of *bla*OXA-58 gene in five *Acinetobacter baumannii* clinical isolates recovered from two hospitals in southern Vietnam during 2012-2014. *A. baumannii* isolates were identified by automated microbiology systems and confirmed by PCR. All isolates were characterized as multidrug resistant by antimicrobials testing using the disk diffusion method. Four imipenem susceptible and one non-susceptible isolates (MIC > 32 *μ*g.ml^−1^) were identified by E-test. PCR amplification of *bla*_OXA-58_ gene upstream and downstream sequences revealed the presence of IS*Aba3* at both locations in one multidrug resistant isolate. Semi quantitation of *bla*_OXA-51_ and *bla*_OXA-58_ gene expression was performed by the 2^−ΔΔCt^ method. The *bla*_OXA-51_ gene expression of five isolates showed little difference but the isolate bearing IS*Aba3-bla*_OXA-58_-IS*Aba3* exhibited significant higher *bla*_OXA-58_ mRNA level. Higher β-lactamases activity in periplasmic than cytoplasmic fraction was found in most isolates. The isolate overexpressing *bla*OXA-58 gene possessed very high periplasmic enzyme activity. In conclusion, the *A. baumannii* isolate bearing IS*Aba3-bla*_OXA-58_ gene exhibited high resistance to imipenem, corresponding to an overexpression of *bla*_OXA-58_ gene and very high periplasmic β-lactamases activity.

## Introduction

Multidrug resistant *A. baumannii* constitutes a serious threat for nosocomial infection control [1]. Carbapenems are currently the antibiotics of choice against multidrug-resistant *Acinetobacter* infections [2] but an increasing rate of resistance to carbapenems was reported worldwide, seriously limiting therapeutic options [3]. Carbapenem-resistant *A. baumannii* has become an alarming health care problem, mainly in developing countries [4]. As a result, carbapenem-resistant *A. baumannii* is classified into the critical priority group according to the urgency of need for new antibiotic treatment and the level of reported antibiotic resistance by World Health Organization [5].

Multiple mechanisms of carbapenem resistance have been identified in *A. baumannii* including low membrane permeability, mutation in its chromosome genes, overexpression of efflux pumps and acquisition of mobile resistance genes [6]. However, the production of carbapenemases is considered as the principal resistance mechanism [7, 8]. The most frequent ones are carbapenem-hydrolyzing class D β-lactamases (CHDLs) and secondly, metalloenzymes (MBL) such as *bla*_NDM_ [9]. In addition, class A β-lactamases such as *bla*_KPC_ gene has recently also detected in *A. baumannii* [10], presenting a serious threat of expanding resistance spectrum in the bacteria.

Currently, six main groups of CHDLs found in *A. baumannii* include *bla*_OXA-51_ -like, *bla*_OXA-23_ -like, *bla*_OXA-24_-like, *bla*_OXA-58_-like genes, *bla*_OXA-143_-like and *bla*_OXA-235_-like [2, 11, 12]. CHDLs exhibit weak carbapenem hydrolysis; however, they can confer resistance mediated by the combination of natural low permeability and IS*Aba* elements located upstream of the gene possibly leading to the gene’s overexpression [2]. Overexpression of *bla*_OXA_ genes usually corresponds to resistance phenotypes [13–15]. Overproduction of oxacillinases, including *bla*_OXA-58_ enzyme, results from the presence of insertion sequences such as IS*Aba1*, IS*Aba2*, IS*Aba3*, or IS*18* which provide strong promoters for gene expression [13, 16].

In Vietnam, *bla*_OXA-23_ is the most widely disseminated class D-carbapenemase in carbapenem-resistant *Acinetobacter baumannii* while *bla*_OXA-24_ is not detected [17]. Even though there is not any information of *bla*_OXA-143_ and *bla*_OXA-235_ in Vietnam up to now, these genes are believed to emerge in other parts of the world [18, 19]. During 2003-2014, the majority of *A. baumannii* clinical isolates recovered harboured *bla*_OXA-51_ and *bla*_OXA-23_ genes. The *bla*_OXA-58_ gene was only detected in isolates recovered from 2010, after the introduction of imipenem in 2008-2009 into hospitals in Vietnam [20, 21]. The *bla*_OXA-58_ - positive isolates investigated in the present study probably emerged at the same time. This recent emergence was in contrast with the striking replacement of *bla*_OXA-58_ by *bla*_OXA-23_ reported in Italy and China for the same period [22, 23]. Furthermore, isolates bearing *bla*_OXA-58_-like gene were recovered from different countries during outbreaks and showed remarkable conserved gene sequence [24–26]. The aim of this study was to investigate genetic structures and relative expression of *bla*_OXA-58_ gene, which lead to imipenem non-susceptibility in clinical isolates recovered from two Vietnamese hospitals during 2012-2014.

## Materials and methods

### Study design

The study focused on *A. baumannii* isolates containing *bla*_OXA-58_ gene with the purpose of determining imipenem-resistance mechanism related to the gene.

### Bacterial isolates, microbial identification and antimicrobial susceptibility testing

Five *A. baumannii* isolates were chosen from a total of 252 non-duplicate *Acinetobacter* spp. isolates recovered from patients admitted to hospitals in southern Vietnam during 2012-2014 and were named DN and TN based on their source hospitals [17]. Microbial isolation and identification in source laboratories were performed using the Phoenix System (BD) and the API 20NE system (bioMerieux). Identification of *A. baumannii* isolates was confirmed by PCR amplification and sequencing of 16S-23S intergenic spacer (ITS) regions. The sequences were deposited in GenBank under accession numbers KY659325, KY659326, KY659327, KY659328 and KY659329. Antimicrobial susceptibility testing was performed by the disk diffusion method and interpreted according to the Clinical and Laboratory Standards Institute guidelines (CLSI, 2014). Tested antimicrobials included ceftazidime, cefotaxime, ceftriaxone, cefpodoxime, cefepime, piperacillin, ampicillin/sulbactam, piperacillin/tazobactam, ticarcillin/clavulanic acid and meropenem, as well as others not belong to β-lactams such as amikacin, gentamicin, ankamycin, netilmicin, ciprofloxacin and levofloxacin. MIC values of imipenem were determined by E-test (bioMérieux); the CLSI-approved breakpoints for imipenem ≥ 8 *μ*g.ml^−1^ and ≤ 2 *μ*g.ml^−1^ were considered as resistant and susceptible, respectively.

### Detection of *bla*_OXA_, *bla*_NDM_ and *bla*_KPC_ genes and insertion sequences

Amplification of *bla*_OXA_ genes including *bla*_OXA-51_, *bla*_OXA-23_, *bla*_OXA-24_ and *bla*_OXA-58_ genes were performed and published in the previous study [21]. *bla*_NDM_ and *bla*_KPC_ genes were amplified in this study as previously reported [27]. The presence of IS*Aba1*, IS*Aba2*, IS*Aba3*, IS*Aba4*, and IS18 was detected as previously described [13, 28]. The sequence of all primers is shown in Table 1.

**Table 1:**
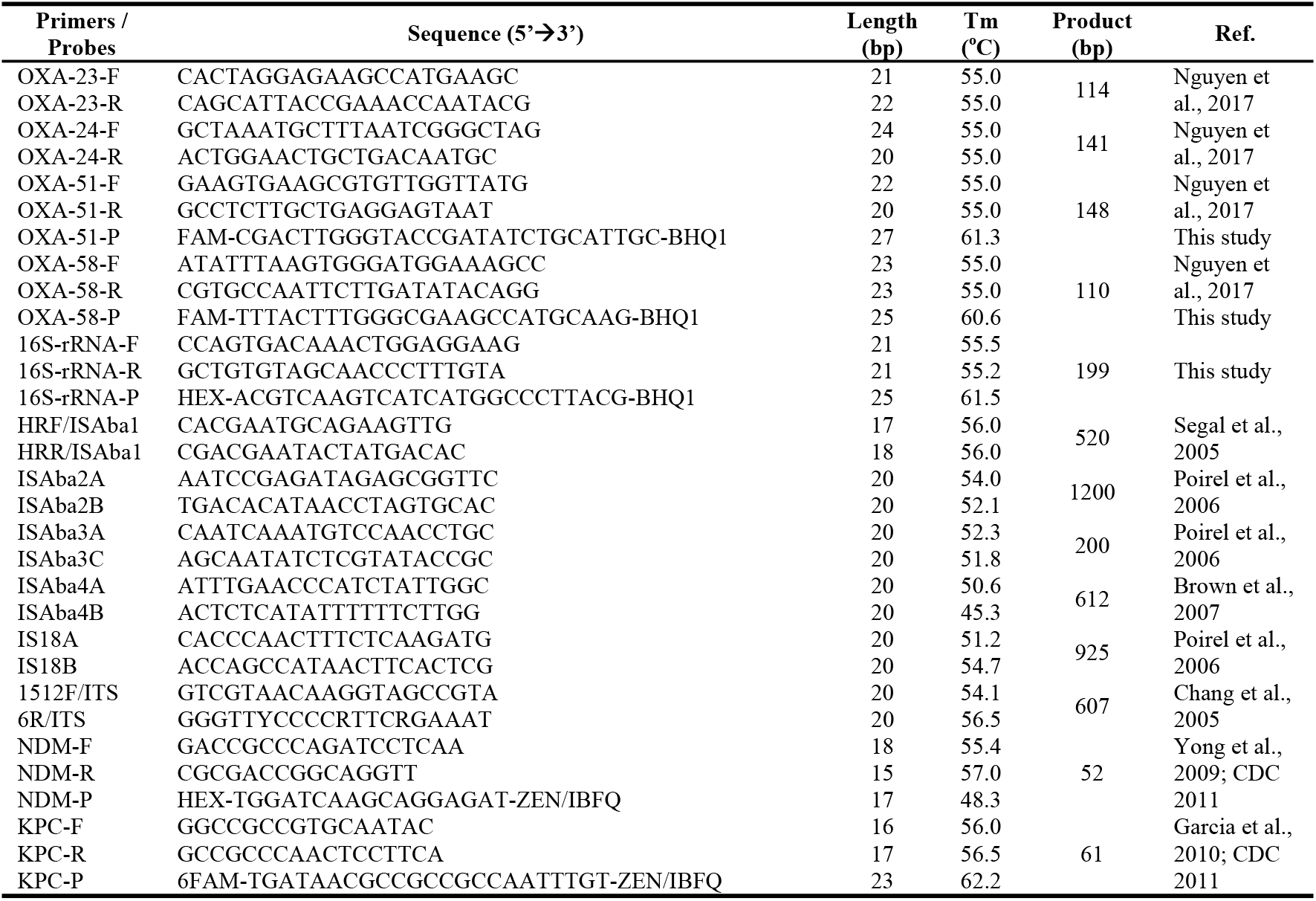
Primers and probes used for PCR amplification and sequencing of antimicrobial resistance genes and related genetic elements

### PCR mapping of *bla*_OXA-58_ and *bla*_OXA-51_ genes

PCR mapping of *bla*OXA genes upstream regions was carried out using combinations of insertion sequence-specific forward primers and *bla*_OXA-51_ and *bla*_OXA-58_ gene-specific reverse primers (Table 1). The presence of IS*Aba3* downstream of *bla*_OXA-58_ was determined by a long-range PCR containing 1X PrimeSTAR GXL Buffer, 0.2 mmol dNTPs, 500 nmol OXA-58-F, 500 nmol ISAba3C, 0.5 U PrimeSTAR GXL DNA polymerase (Takara). PCR products were sent to 1-BASE (https://order.base-asia.com/) for purification and sequencing. The sequences were analysed by BioEdit 7.0.9.0. (http://www.mbio.ncsu.edu/BioEdit/bioedit.html) and sequence similarity was assessed using the BLAST program (https://blast.ncbi.nlm.nih.gov/Blast.cgi). The sequence of *bla*OXA-58 and its surrounding IS*Aba3* were deposited in GenBank under accession number KY660721.

### Analysis of *bla*_OXA-58_ and *bla*_OXA-51_ gene expression by real-time RT-PCR

Mid-log phase of bacterial cultures was treated with 1 *μ*mol.ml^−1^ oxacillin for 24 h and was subsequently used for RNA extraction [29]. Treatment with RNAse-free DNAse I (Sigma) was performed at 37°C for 2 h. The concentration and DNase-free quality of RNA samples were spectrophotometrically assessed and confirmed by the amplification of chromosomal *bla*_OXA-51_ and 16S rRNA. Fifteen microliters of each RNA sample were reverse-transcribed in a final volume of 20 *μ*l containing random hexamers, MMLV reverse transcriptase (Agilent) at 42° C for 45 min.

Amplification of *bla*_OXA-51_, *bla*_OXA-58_, and 16S rRNA was performed in a final volume of 25 μl containing 5 *μ*l cDNA, 3 mmol MgCl_2_, 200 nmol dNTPs, 2 U h-Taq DNA polymerase (Solgent), 300 nmol of OXA-51/58-F/R primers, 150 nmol of OXA-51/58-P probes, 200 nmol of 16S-F/R primers, 100 nmol of 16S-P probe (IDT). Primer and probe sequences were given in Table 1. Each real-time PCR was performed in triplicate on the Stratagene Mx3005P real-time PCR system (Agilent). The reaction mixture was incubated for 15 min at 95°C, followed by 40 cycles of 10 s at 95°C and 20 s at 60°C. Normalized expression of *bla*_OXA-51_ and *bla*_OXA-58_ genes was calculated relatively to the 16S rRNA reference gene according to the 2^−ΔΔCt^ method [30].

### Multiple-locus variable number tandem repeat analysis

Multiple-locus variable number tandem repeat analysis (MLVA) as previously described [17, 31, 32] was used to profiling the *A. baumannii* isolates in the study. The method works on eight variable number tandem repeats (VNTR) loci naming 3468, 1988, 3002, 845, 2396, 5350, 826, and 2240 to determine relatedness among the *A. baumannii* isolates.

### β-lactamases extraction and quantitation

Isolates were grown on LB medium supplemented with 1 *μ*mol.ml^−1^ oxacillin for 18-24 h at 37° C in a shaking incubator. The supernatants (extracellular fraction) were collected after centrifugation of bacterial cultures and precipitated with absolute ethanol (1:4) in 20 min at −20° C [33]. Periplasmic fractions were recovered from cell pellets [34]. Protein concentration was determined by the Bradford method [35].

β-lactamases activity was determined based on nitrocefin hydrolysis [36, 37]. Briefly, 1-5 *μ*l extracellular and periplasmic fractions obtained from each isolate were incubated with 40 nmol nitrocefin dissolved in 0.1M phosphate buffer, pH 7.0 in a total volume of 100 *μ*l. Samples were loaded onto microtiter plates and the absorbance at 482 nm was measured kinetically at room temperature for 2-30 minutes using an ELISA spectrophotometer. The specific β-lactamase activity was calculated and expressed as mU.mg^−1^ of protein based on the quotient of β-lactamase activity (mU.ml^−1^) and protein concentration (mg.ml^−1^).

### Statistical analysis

Analysis of variance (ANOVA) was used to analyse the difference among β-lactamase activity means of isolates. t-test was used to determine the significant difference of extracellular and periplasmic β-lactamase activity. A *P* value <0.05 was considered significant.

## Results and Discussion

*bla*_OXAs_ are prevalent in *A. baumannii*. We had previously performed *bla*_OXAs_ identification in *A. baumannii* isolates from three hospitals in Southern Vietnam and found *bla*_OXA-23_ was dominant [17]. Even though *bla*_OXA-58_ existed with a small number in Vietnamese population, the exact genetic context involving antimicrobial resistance elements remained unknown. Here, we uncovered the imipenem resistance mechanism of *bla*_OXA-58_-positive *A. baumannii* isolates. The overexpression of *bla*_OXA-58_ gene has been seen in the isolate with high resistance phenotype through relative quantification of mRNA of the corresponding gene. The specific possible-intact IS*Aba3* sequence upstream of *bla*_OXA-58_ gene could be the key factor for the high expression. In addition, the high β-lactamase activity in the periplasmic space observed in the study could be the outcome of the phenomenon.

### Antimicrobial susceptibility testing

All five isolates (DN050, TN078, DN014, TN341, TN345) were classified as multidrug resistant (MDR) since they were non-susceptible to at least one agent in three or more antimicrobial categories including aminoglycosides, antipseudomonal carbpenems, antipseudomonal fluoroquinolones, antipseudomonal penicillins and β-lactamase inhibitors, extended-spectrum cephalosporins, folate pathway inhibitors, penicillins and β-lactamase inhibitors, polymyxins, and tetracyclines [38]. In this study, although several antimicrobials were not tested because of the availableness of them at different times and hospitals, all isolates satisfied the definition to be defined as MDR. Isolate DNA050 was non-susceptible to all antimicrobials tested. The other four was all susceptible to imipenem (there were three isolates non-susceptible to meropenem as hospitals reported), but for other antimicrobials, their susceptibility varied. Isolate TN078 and DN014 were non-susceptible to three categories while isolate TN341 and TN345 were non-susceptible to five categories (Table 2).

**Table 2:** β-lactams susceptibility profiles and genotypes of five *A. baumannii* isolates

| Isolate | ID | DN050 | TN078 | DN014 | TN341 | TN345 |
| --- | --- | --- | --- | --- | --- | --- |
| Phenotype | Imipenem | + | - | - | - | - |
|  | Meropenem | + | N/A | + | + | + |
|  | Ceftazidime | + | + | - | + | + |
|  | Cefotaxime | + | + | + | N/A | + |
|  | Ceftriaxone | + | + | + | + | + |
|  | Cefpodoxime | N/A | + | N/A | + | + |
|  | Cefepime | + | - | + | + | + |
|  | Piperacillin | + | N/A | - | N/A | N/A |
|  | Ampicillin/Sulbactam | + | - | - | + | + |
|  | Piperacillin/Tazobactam | + | - | - | + | + |
|  | Ticarcillin/Clavulanic acid | N/A | + | N/A | + | + |
|  | Amikacin | + | + | N/A | N/A | N/A |
|  | Gentamicin | + | + | N/A | + | + |
|  | Ankamycin | N/A | N/A | + | + | - |
|  | Netilmicin | N/A | N/A | - | + | + |
|  | Ciprofloxacin | + | - | + | + | + |
|  | Levofloxacin | + | - | - | N/A | N/A |
| Genotype | *bla<sub>OXA-51</sub>* | + | + | + | + | + |
|  | *bla<sub>OXA-23</sub>* | - | - | - | - | - |
|  | *bla<sub>OXA-24</sub>* | - | - | - | - | - |
|  | *bla<sub>OXA-58</sub>* | + | + | + | + | + |
|  | *ISAba1* | + | + | - | + | - |
|  | *ISAba2* | + | + | + | - | - |
|  | *ISAba3* | + | + | + | + | + |
|  | *ISAba4* | - | - | - | - | - |
|  | *IS18* | - | - | - | - | - |
|  | *ISAba3_bla<sub>OXA-58</sub>* | + | - | - | - | - |
|  | *NDM* | + | + | - | - | - |
|  | *KPC* | - | - | - | - | - |
|  | MLVA profile* | 6-0-7-1-17-5-0-3 | 6-0-7-14-17-6-15-3 | 9-0-7-1-7-5-14-3 | 9-0-5-15-15-6-0-2 | 9-0-5-15-15-6-0-2 |

N/A, not determined; -, assay negative (susceptible/absence); +, assay positive (resistant/presence)
\* MLVA profile according to the surveyed loci: 3468-1988-3002-845-2396-5350-826-2240

### Isolates genotyping and profiling

All isolates were identified as *A. baumannii* based on 16S-23S intergenic spacer (ITS) regions sequencing. Based on MLVA profiling, four different MLVA types within the five isolates reflected substantial genetic diversity in the sampled Vietnamese *A. baumannii* isolates, as previously described [17].

No isolate with *bla*_KPC_ gene was detected, while two isolates contained *bla*_NDM_ gene (DN050 and TN078). Even though the two isolates were singletons (based on MLVA types from previous study [17]) with different phenotypes, they had close relatedness with just difference in 3/8 loci surveyed and very similar resistance determinants, especially the *bla*_NDM_ gene. Therefore, the difference in resistance phenotype was mostly because of the distinguished genotype with ISAba3_*bla*_OXA-58_ in isolate DN050, compared to isolate TN078. It might be necessary for *bla*_NDM_ gene locating in a specific genetic context to be expressed as one of the important and strong resistance determinants. The mechanism should be explored further.

Regardless of the genetic diversity of the isolates, the *bla*_OXA-58_ gene sequence analysis (data not shown) of all isolates was identical with the reported *bla*_OXA-58_ gene [39]. This was in agreement with previous work showing a lack of diversity in this gene, probably due to its recent acquisition by *A. baumanii* from other species [3].

All isolates were *bla*_OXA-58_ and *bla*_OXA-51_ positive, *bla*_OXA-23_ and *bla*_OXA-24_ negative (Table 2). Analysis of insertion sequences revealed the presence of IS*Aba1* and IS*Aba2* but they were not located upstream of *bla*_OXA-51_ nor *bla*_OXA-58_ genes in all isolates. IS*Aba4* and IS18 were not detected. IS*Aba3* was detected in all isolates (Table 2). However, only isolate DN050 possessed a *bla*_OXA-58_ gene bracketed by two IS*Aba3* elements (Figure 1, Figure 2). The promoter region of *bla*_OXA-58_ gene in this isolate (Figure 2) was similar to sequences described by Poirel et al. [39]. The genetic structure of *bla*_OXA-58_ upstream sequences which led to overexpression of this gene displayed a remarkable variability [39–41]. Hybrid promoters constituted of an IS*Aba3* sequence truncated by other insertion sequences were generally considered as strong promoters [23, 42]. However, in this study isolate DN050 bearing possible-intact IS*Aba3* sequence upstream of *bla*_OXA-58_ gene was not interrupted by inserted sequences, provided −35 and −10 promoter sequences as already described [39]. This structure probably drove high level carbapenemase production. The acquisition of insertion sequences by an imipenem-susceptible *bla*_OXA−58_ harboring isolate can lead to carbapenem resistance in *A. baumannii* [39]. Our results highlighted the threat of undetected reservoirs of carbapenem resistant determinants and mechanisms in Vietnamese *A. baumannii* isolates.

**Figure 1:**
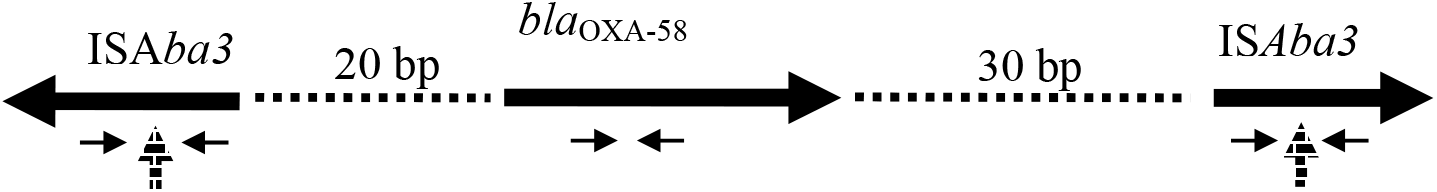
Genetic structures identified in the IS*Aba3-bla*_OXA-58_ positive *A. baumannii* isolate, DN050. IS*Aba3* and *bla*_OXA-58_ gene were indicated by horizontal bold arrows. Horizontal dash lines indicated sequences separating IS*Aba3* and *bla*_OXA-58_. Vertical arrows were for the truncated sites previously reported that did not exist in this isolate. Positions of primers were indicated as referred to Table 1 with short thin arrows. The figure is not to scale.

**Figure 2:**
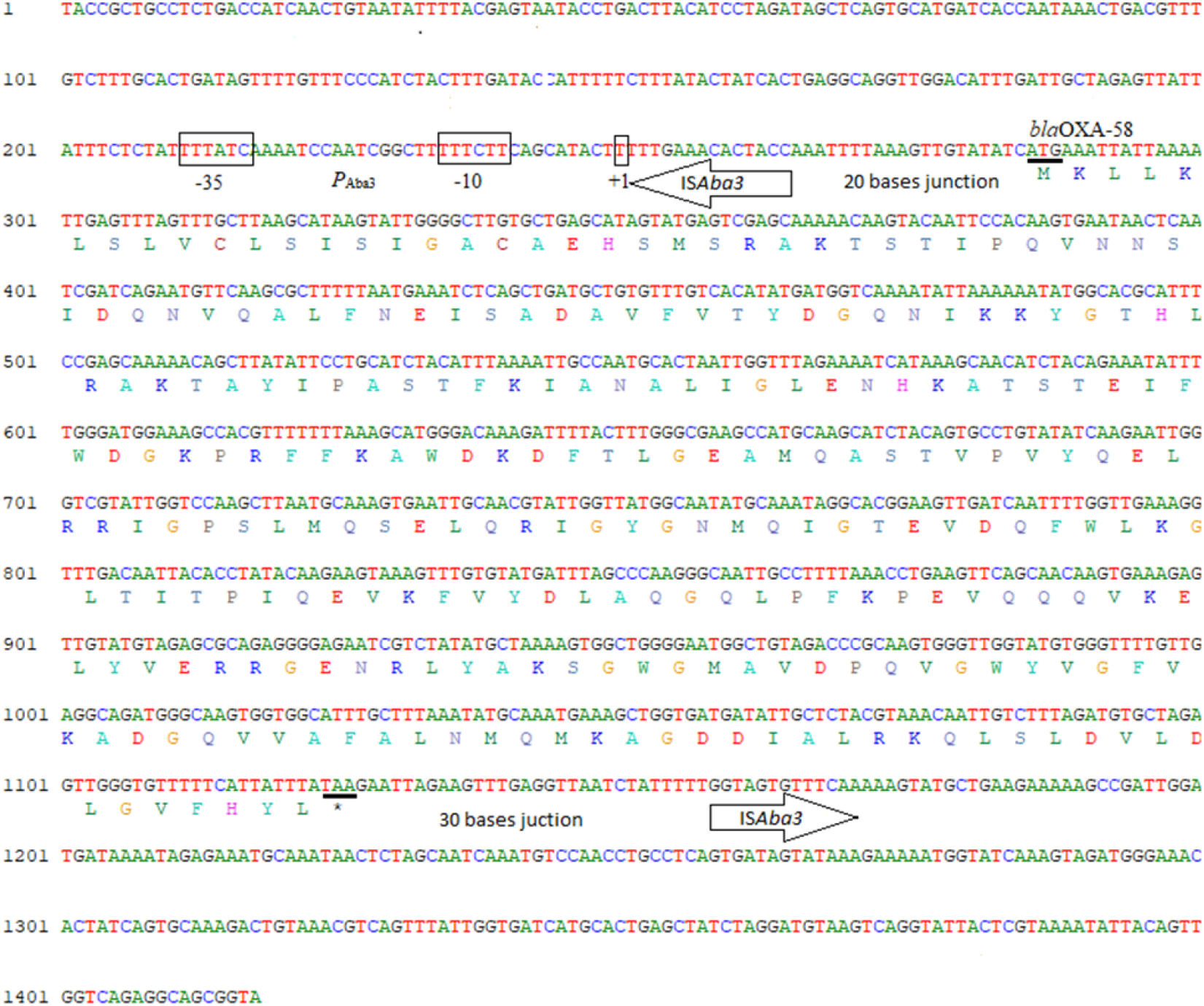
Promoter structure of *bla*_OXA-58_ gene from isolate DN050. The −35 and −10 putative promoter sequences and the +1 transcription initiation site within IS*Aba3* are boxed. The *bla*_OXA-58_ start and stop codons, ATG (M) and TAA (*) respectively, are underlined. Upstream IS*Aba3*/*bla*_OXA-58_ sequences and downstream IS*Aba3*/*bla*_OXA-58_ gene junctions are indicated by arrows. Full sequences obtained are deposited in GenBank (accession number KY660721).

### Relative quantitation of *bla*_OXA-58_ and *bla*_OXA-51_ mRNA level

We chose three isolates (DN050, TN341 and TN345) to study the relative expression of *bla*_OXA-51_ and *bla*_OXA-58_ under condition with oxacillin as an inducer and without oxacillin induction. They all had high β-lactamase activity in periplasmic fractions as shown in the experiment followed (Table 3). The mRNA level of *bla*_OXA-58_ and *bla*_OXA-51_ gene in all isolates was determined by quantitative real-time RT-PCR. Under oxacillin induction, DN050 showed a significantly higher level of *bla*_OXA-58_ mRNA expression than isolates TN341 and TN345 (Figure 3). *bla*_OXA-51_ expression was also upregulated, but not comparable to that of *bla*_OXA-58_. Interestingly, the high expression level of *bla*_OXA-58_ from DN050 could be associated with the presence of an upstream IS*Aba3* sequence as previously suggested [39]. Furthermore, in this study the possible-intact IS*Aba3* sequence might be customized to *bla*_OXA-58_ gene to drive a very strong gene expression, as seen in IS*Aba1* for *bla*_OXA-23_ and AmpC genes [43]. The other isolates lacked upstream IS*Aba3* sequence.

**Table 3:** Relative quantitation of *bla*_OXA-51_ and *bla*_OXA-58_ mRNA level and β-lactamase activity in five *A. baumannii* isolates

| Isolate |  | DN050 | TN078 | DN014 | TN341 | TN345 |
| --- | --- | --- | --- | --- | --- | --- |
| MIC imipenem ($\mu\text{g.ml}^{-1}$) | | $\geq 32$ | 0.5 | 0.19 | 0.75 | 0.5 |
| Relative expression of *bla*<sub>OXA-51</sub> | $\Delta\text{Ct}$ | $8.87 \pm 0.39$ | $12.87 \pm 0.30$ | $7.54 \pm 0.66$ | $9.64 \pm 0.71$ | $8.80 \pm 0.60$ |
|  | Expression (time) | 1.00<br>(0.76-1.31) | 0.06<br>(0.05-0.08) | 2.51<br>(1.58-3.98) | 0.59<br>(0.36-0.96) | 1.04<br>(0.69-1.59) |
| Relative expression of *bla*<sub>OXA-58</sub> | $\Delta\text{Ct}$ | $4.18 \pm 1.18$ | $8.64 \pm 0.48$ | $7.99 \pm 2.23$ | $8.45 \pm 0.53$ | $7.28 \pm 0.24$ |
|  | Expression (time) | 25.69<br>(11.3-58.37) | 1.17<br>(0.84-1.64) | 1.84<br>(0.39-8.61) | 1.33<br>(0.93-1.93) | 3.01<br>(2.55-3.56) |
| Total β-lactamase activity ($\text{U.mg}^{-1}$) | Extracellular | $10.8 \pm 3.3$ | $17.7 \pm 5.2$ | $13.9 \pm 4.1$ | $10.8 \pm 3.3$ | $12.9 \pm 3.8$ |
| | Periplasmic | $44.7 \pm 12.8$ | $15.3 \pm 4.3$ | $28.5 \pm 8.2$ | $49.6 \pm 16.0$ | $27.4 \pm 10.0$ |

**Figure 3:**
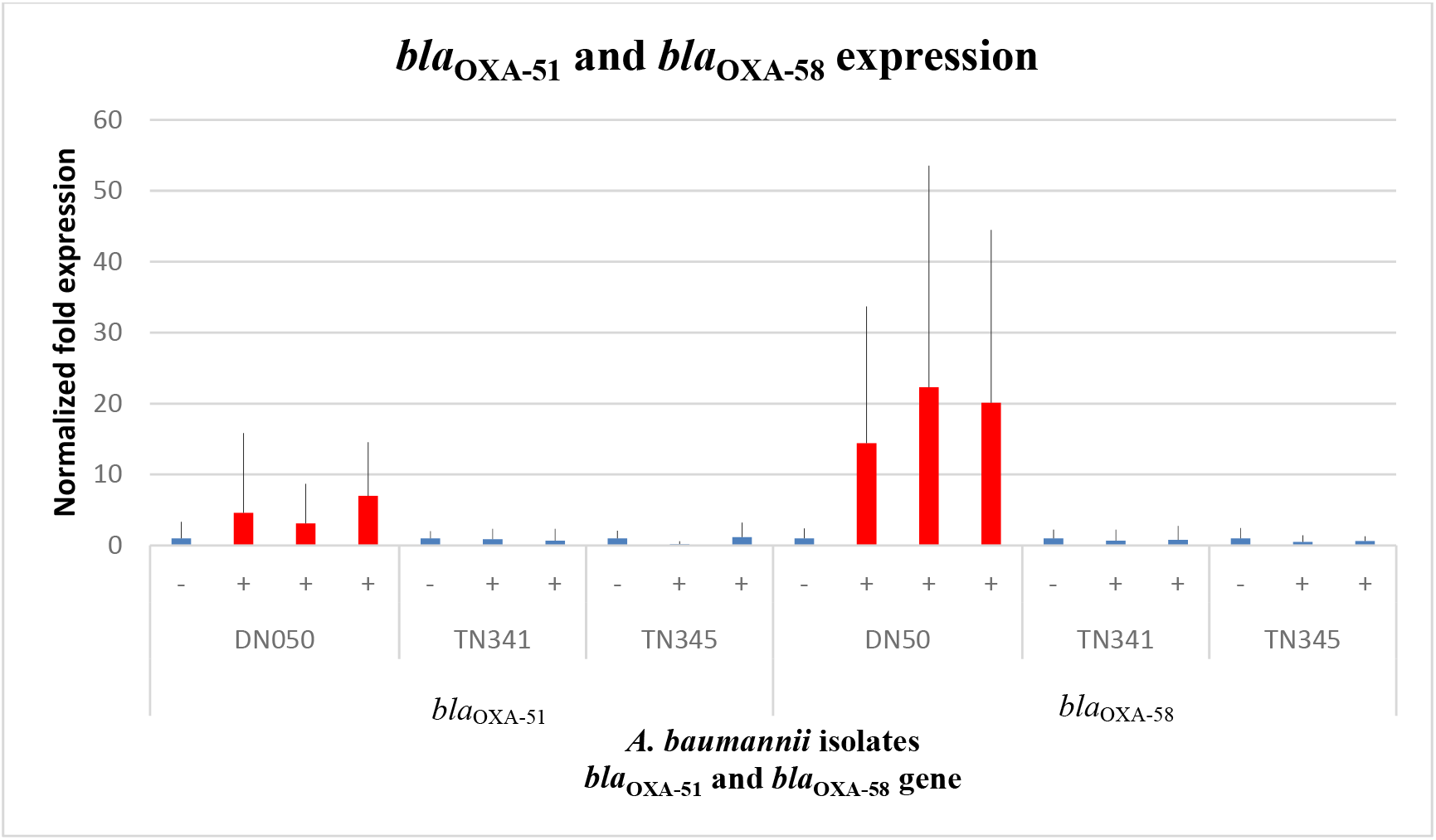
Duplex real-time RT-PCR analysis of the *bla*_OXA-51_ and *bla*_OXA-58_ mRNA relative expression in three *A. baumannii* isolates. The error bars represent the deviation for the normalized fold expression of *bla*_OXA-51_ and *bla*_OXA-58_ in three isolates which were positive or negative for the IS*Aba3* upstream of the *bla*_OXA-58_ gene. (−), not induced; (+), induced

### Analysis of periplasmic β-lactamases activity in association with *bla*_OXA-51/-58_-relative expression

Under condition of oxacillin induction, the *bla*_OXA-58_ expression of isolate DN050 (MIC_imipenem_ ≥ 32 μg.ml^−1^) was also significantly higher than the expression of other four isolates, TN078, DN014, TNA341 and TN345 with MIC_imipenem_ were 0.5, 0.19, 0.75 and 0.5 respectively (Table 3). All isolates expressed low level of *bla*_OXA-51_, confirming that the presence of *bla*_OXA-51_, without an upstream IS*Aba1*, did not confer a resistance phenotype [16]. Furthermore, in variants harbouring *bla*_OXA-51_ and *bla*_OXA-58_ genes, carbapenem resistance only correlated with *bla*_OXA-58_ [44], which is in agreement with the results of this study.

The enzyme activity of extracellular fractions was not significantly different (*p* = 0.2187) while the one of periplasmic fractions exhibited significant difference among isolates (*p* < 0.0001). Extracellular fractions possessed lower enzyme activity than periplasmic fractions (*p*=0.0355) in most cases. The periplasmic fraction recovered from all isolates exhibited variable β-lactamases activity, with very high activity corresponding to isolate DN050. Isolates TN341 displayed the highest β-lactamase activity though weakly expressed *bla*_OXA-58_ gene. This high enzyme activity probably corresponded to other β-lactamases responsible for the multidrug resistance phenotypes of the isolate, such as extended-spectrum AmpCs [45]. The presence of other β-lactamases could explain the high enzyme activity in periplasmic fractions of the other isolates. Particularly, *bla*_NDM_ gene detected in both isolate DN050 and TN078, but the corresponding β-lactamase activities as well as the antimicrobial susceptibilities were different between the two isolates. The mechanism that a strain carrying a *bla*_NDM_ gene is not resistant to carbapenems needs to be discovered further in *A. baumannii*. It might need a unique genetic structure for *bla*_NDM_ gene to be expressed as seen in *K. pneumoniae* [46].

In a transformed *A. baumannii* strain with a *bla*_OXA-58_ plasmid-borne vector, this carbapenemase is selectively released via outer membrane vesicles (OMV) after periplasmic translocation through Sec-dependent system [33]. Furthermore, overexpression of *bla*_OXA-58_ gene increases its periplasmic enzyme concentration and extracellular release leading to efficient carbapenem hydrolysis [33]. The *bla*_OXA-58_ high mRNA level and high periplasmic β-lactamase activity of the DN050 isolate in this study suggested a similar overexpression, periplasmic translocation and release mechanism of *bla*_OXA-58_ carbapenemase, even though our experimental work did not directly show the selection of OMV after being translocated to periplasmic space. The high periplasmic β-lactamases activity of the isolates, especially TN341 in this study also suggested a possible translocation and release of other β-lactamases with a mechanism similar to that identified with *bla*_OXA-58_. Further studies should be carried out to prove the suggested mechanism in clinical isolate similar to the transformed *A. baumannii* strain. To the best of our knowledge, our study is the first report on the overexpression of *bla*_OXA-58_ gene of *A. baumannii* clinical isolates from Vietnam.

This study had some limitations. The first limitation involved the small sample size due to the low prevalence of clinical isolates harbouring *bla*_OXA-58_ gene in the population surveyed. The screening has been done in previous studies [17]. Secondly, we did not characterize other resistance mechanisms in *A. baumannii* such as the overexpression of efflux pump genes or existence of multicopy *bla*_OXA-58_ gene [7, 11, 47]. In addition, the presence of other β-lactamase genes such as *bla*_IMP_, *bla*_VIM_, *bla*_GES_, *bla*_OXA-143_ and *bla*_OXA-_ 235 was not excluded. Furthermore, we did not carry out an alternative experimental approach, such as western blotting against *bla*_OXA-58_ to unequivocally determine if the increase in β-lactamase activity is mainly due to this protein.

## Conclusions

This study identified a mechanism of imipenem resistance related to overexpression of *bla*_OXA-58_ gene preceded by IS*Aba3* and its corresponding periplasmic enzyme present at high concentration in a multidrug resistant clinical isolate recovered from a hospital in Vietnam.

## Data availability

Data used to support the findings of this study are available from the corresponding author upon request.

## Acknowledgements

We thank Dao Minh Y from Dong Nai general hospital and Nguyen Si Tuan from Thong Nhat – Dong Nai general hospital for providing clinical *A. baumannii* isolates used in this study.

## Supplementary Materials

Related information of E-test images, electrophoresis results of PCR screening for the presence / absence of ISs, duplex real-time RT-PCR data analysis, Bradford assay standard curve and Nitrocefin standard curve for protein quantification and β-lactamase activity determination of supernatant and periplasmic fractions were detailed in Supplementary Materials.

