## Supplemental file for "Overexpression of *bla*_OXA-58_ Gene Driven by IS*Aba3* is Associated with Imipenem Resistance in a Clinical *Acinetobacter baumannii* Isolate from Vietnam"

### Supplementary Material

#### Tables of contents


- Figure S1. Result of imipenem E-test for five clinical isolates of *A. baumannii* (*bla*$_{OXA-58}$)
- Figure S2. Electrophoresis results of PCR screening for the presence / absence of IS*Aba1*, IS*Aba2*, IS*Aba3*, IS*Aba4*, and IS18 in five clinical isolates of *A. baumannii* (*bla*$_{OXA-58}$)
- Figure S3. Electrophoresis results of PCR for the presence / absence of insertion sequence (IS) upstream of *bla*$_{OXA-58}$ gene.
- Figure S4. Duplex real-time RT-PCR analysis of the *bla*$_{OXA-51}$ and *bla*$_{OXA-58}$ mRNA relative expression compared with 16S rRNA in five *A. baumannii* isolates.
- Figure S5. Bradford assay standard curve of concentration versus absorbance for protein quantification.
- Figure S6. Nitrocefin Standard Curve.
- Table S1. Duplex real-time RT-PCR analysis of the *bla*$_{OXA-51}$ and *bla*$_{OXA-58}$ mRNA relative expression in three *A. baumannii* isolates under conditions with oxacillin as an inducer or without oxacillin induction.
- Table S2. Results for protein quantification of supernatant and periplasmic fractions.
- Table S3. Results for β-lactamase activity of supernatant and periplasmic fractions.

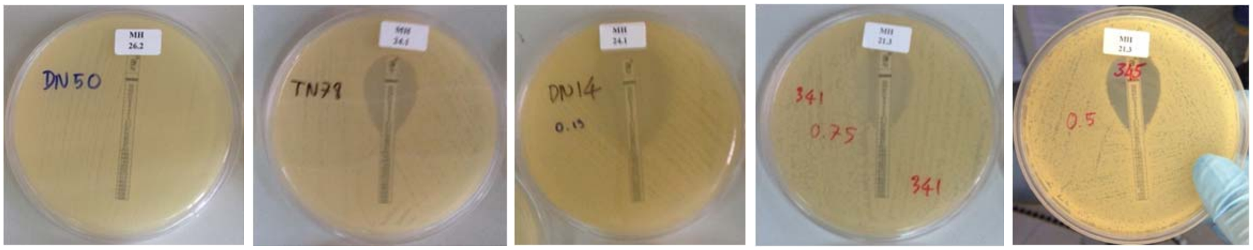


Figure S1. Result of imipenem E-test for five clinical isolates of *A. baumannii* (*bla*$_{OXA-58}$)

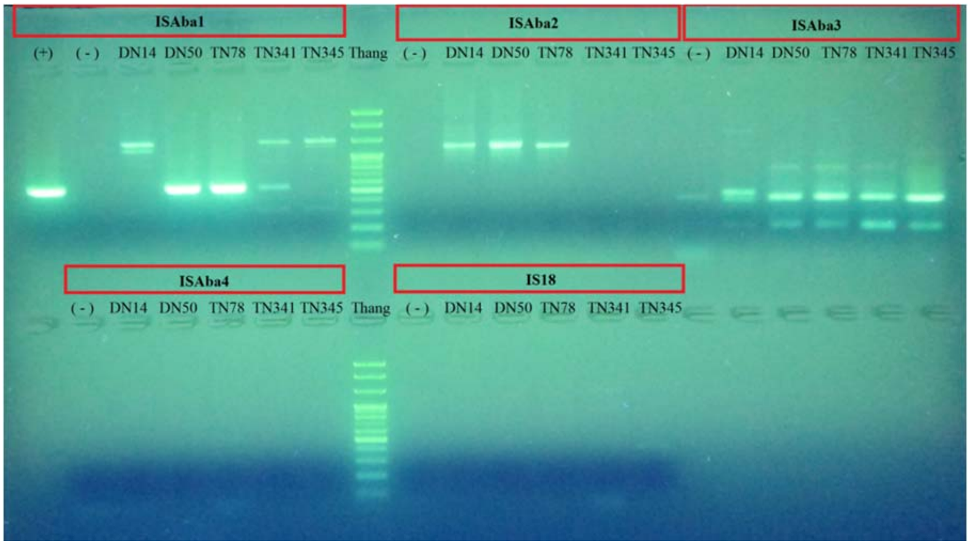


Figure S2. Electrophoresis results of PCR screening for the presence / absence of IS*Aba1*, IS*Aba2*, IS*Aba3*, IS*Aba4*, and IS18 in five clinical isolates of *A. baumannii* (*bla*$_{OXA-58}$)

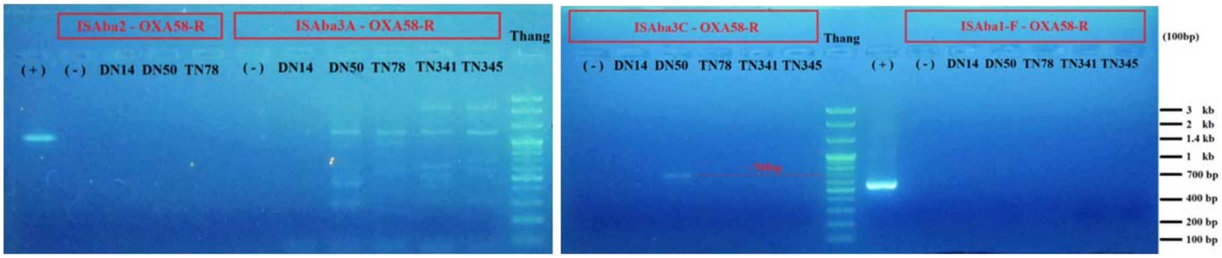


Figure S3. Electrophoresis results of PCR for the presence / absence of insertion sequence (IS) upstream of *bla*$_{OXA-58}$ gene.

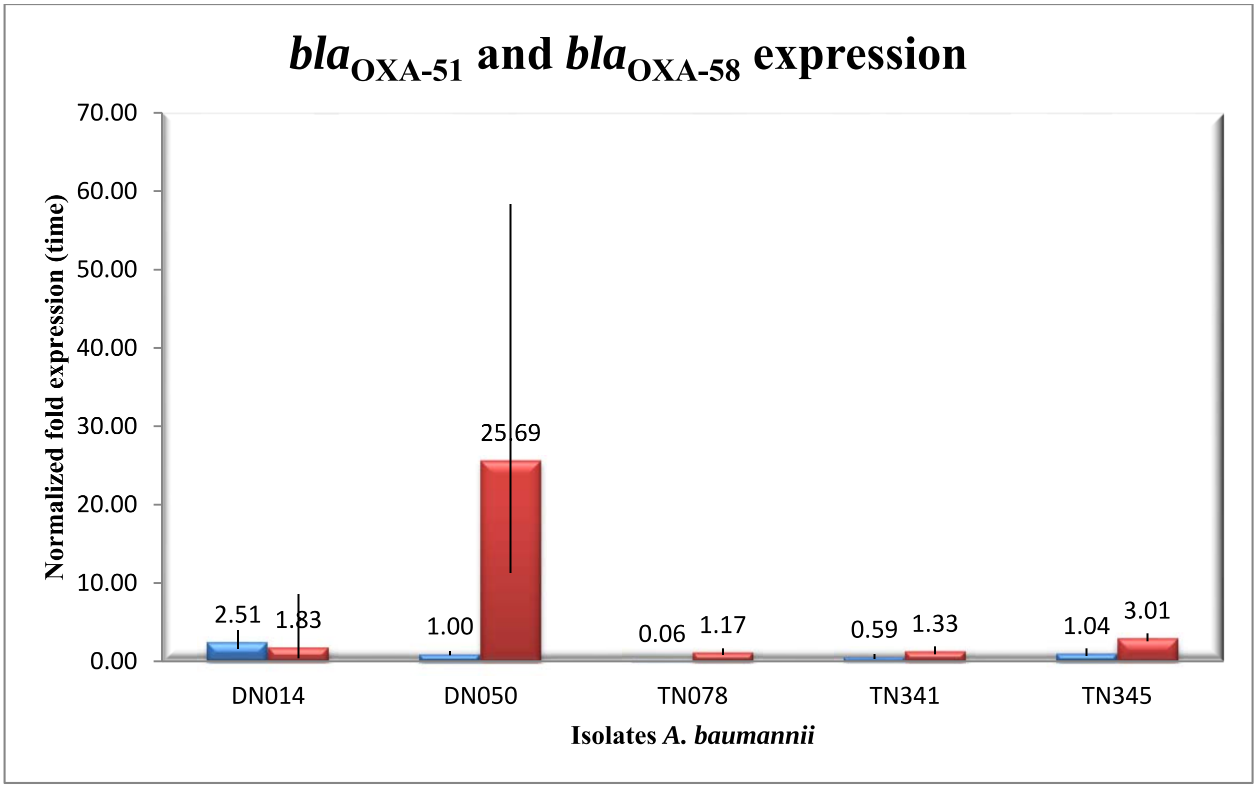


Figure S4. Duplex real-time RT-PCR analysis of the *bla*$_{OXA-51}$ and *bla*$_{OXA-58}$ mRNA relative expression normalized to 16S rRNA gene as a reference in five *A. baumannii* isolates. The expression of *bla*$_{OXA-51}$ in isolate DN050 is used as a calibrator. Error bars represent deviation for the normalized fold expressions of *bla*$_{OXA-51}$ and *bla*$_{OXA-58}$ in five isolates induced by oxacillin. Blue and red colours represent the expression of *bla*$_{OXA-51}$ and *bla*$_{OXA-58}$ correspondingly.

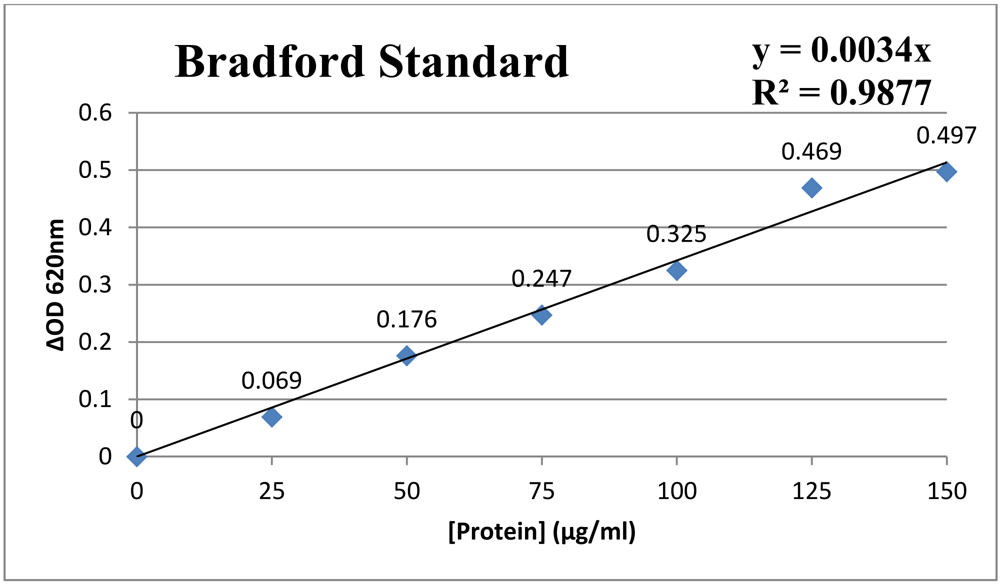


Figure S5. Bradford assay standard curve of concentration versus absorbance for protein quantification.

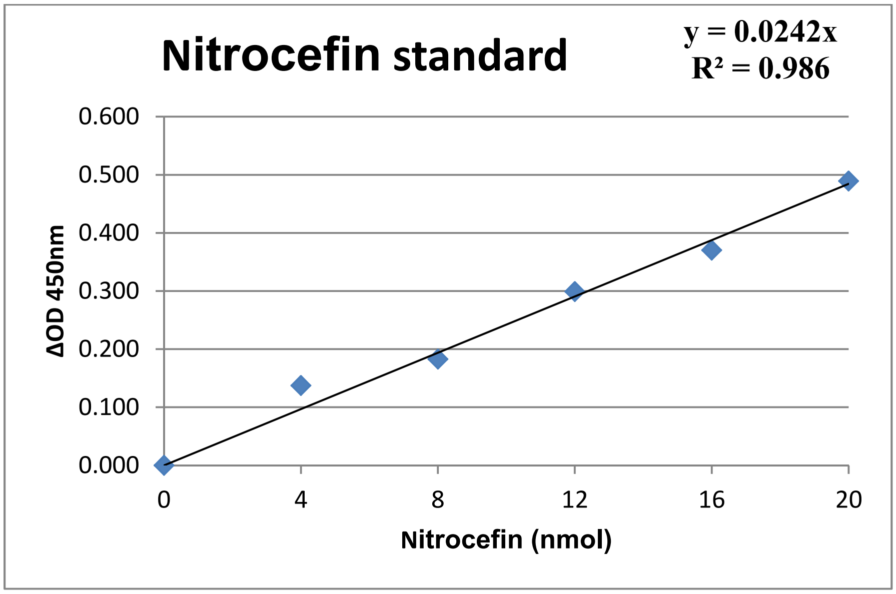


Figure S6. Nitrocefin Standard Curve

Table S1. Duplex real-time RT-PCR analysis of the *bla*$_{OXA-51}$ and *bla*$_{OXA-58}$ mRNA relative expression in three *A. baumannii* isolates under conditions with oxacillin as an inducer or without oxacillin induction. 16S rRNA gene is used as the reference for normalization and the non-induced control is run as the calibrator.

| **Isolate** | | **DN050** | | | **TN341** | | | **TN345** | | |
|---|---|---|---|---|---|---|---|---|---|---|
| | | Min | Value | Max | Min | Value | Max | Min | Value | Max |
| Relative expression of *bla*$_{OXA-51}$ (time) | Not induced | 0.4 | 1.0 | 2.3 | 1.0 | 1.0 | 1.0 | 0.9 | 1.0 | 1.1 |
| | Induced 1 | 1.9 | 4.60 | 11.2 | 0.5 | 0.9 | 1.5 | 0.1 | 0.2 | 0.4 |
| | Induced 2 | 1.8 | 3.10 | 5.6 | 0.3 | 0.7 | 1.6 | 0.8 | 1.2 | 2.0 |
| | Induced 3 | 6.5 | 7.0 | 7.6 | - | - | - | - | - | - |
| Relative expression of *bla*$_{OXA-58}$ (time) | Not induced | 0.7 | 1.0 | 1.4 | 0.8 | 1.0 | 1.2 | 0.7 | 1.0 | 1.5 |
| | Induced 1 | 10.7 | 14.4 | 19.3 | 0.3 | 0.7 | 1.5 | 0.3 | 0.5 | 0.9 |
| | Induced 2 | 15.9 | 22.3 | 31.3 | 0.3 | 0.8 | 2.0 | 0.5 | 0.6 | 0.7 |
| | Induced 3 | 16.6 | 20.1 | 24.4 | - | - | - | - | - | - |

Table S2. Results for protein quantification of supernatant and periplasmic fractions. The concentration of protein (µg/ml) was determined using the equation y = 0.0034x with an $R^2$ value of 0.9877, where y is absorbance and x is concentration.

| **Isolate** | **DN050** | **TN078** | **DN014** | **TN341** | **TN345** |
|---|---|---|---|---|---|
| Supernatant (µg/ml) | 58.82 | 45.29 | 37.06 | 47.94 | 45.88 |
| Periplasmic (µg/ml) | 22.94 | 85.00 | 26.47 | 18.24 | 37.35 |

Table S3. Results for β-lactamase activity of supernatant and periplasmic fractions. The hydrolyzed Nitrocefin (nmol) generated by β-lactamase during the reaction time (ΔT) was determined using the equation y = 0.0242x with an $R^2$ value of 0.986, where y is absorbance and x is the amount of hydrolyzed Nitrocefin. The β-lactamase activity of the test samples is calculated based on the formula: B/(ΔTxV)xD (nmol/min/ml) or (mU/ml) where B is the amount of Nitrocefin from the Standard Curve (nmol), ΔT is the reaction time (min), V is the sample volume added into the reaction well (ml), D is the sample dilution factor.

| **Sample** | **Isolate** | **ΔOD** | **ΔT (min)** | **B (nmol)** | **V (ml)** | **Diluted (D)** | **β-lactamase activity (mU/ml)** | **β-lactamase activity (mU/µg)** | **SD** |
|---|---|---|---|---|---|---|---|---|---|
| **Supernatant** | DN050 | 0.385 | 25 | 15.9 | 0.005 | 5 | 636.4 | 10.8 | 3.3 |
| | TN078 | 0.388 | 20 | 16.0 | 0.005 | 5 | 801.7 | 17.7 | 5.2 |
| | DN014 | 0.299 | 24 | 12.4 | 0.005 | 5 | 514.8 | 13.9 | 4.1 |
| | TN341 | 0.188 | 15 | 7.8 | 0.005 | 5 | 517.9 | 10.8 | 3.3 |
| | TN345 | 0.201 | 14 | 8.3 | 0.005 | 5 | 593.3 | 12.9 | 3.8 |
| **Periplasmic** | DN050 | 0.310 | 25 | 12.8 | 0.005 | 10 | 1024.8 | 44.7 | 12.8 |
| | TN078 | 0.251 | 16 | 10.4 | 0.005 | 10 | 1296.5 | 15.3 | 4.3 |
| | DN014 | 0.219 | 24 | 9.0 | 0.005 | 10 | 754.1 | 28.5 | 8.2 |
| | TN341 | 0.197 | 18 | 8.1 | 0.005 | 10 | 904.5 | 49.6 | 16.0 |
| | TN345 | 0.124 | 10 | 5.1 | 0.005 | 10 | 1024.8 | 27.4 | 10.1 |
